# Singular manifolds of proteomic drivers to model the evolution of inflammatory bowel disease status

**DOI:** 10.1101/751289

**Authors:** Ian Morilla, Thibaut Léger, Assiya Marah, Isabelle Pic, Hatem Zaag, Eric Ogier-Denis

## Abstract

The conditions who denotes the presence of an immune disease are often represented by interaction graphs. These informative, but complex structures are susceptible to being perturbed at different levels. The mode in which that perturbation occurs is still of utmost importance in areas such as reprogramming therapeutics. In this sense, the overall graph architecture is well characterise by module identification. Topological overlap-related measures make possible the localisation of highly specific module regulators that can perturb other nodes, potentially causing the entire system to change behaviour or collapse. We provide a geometric framework explaining such situations in the context of inflammatory bowel diseases (IBD). IBD are important chronic disorders of the gastrointestinal tract which incidence is dramatically increasing worldwide. Our approach models different IBD status as Riemannian manifolds defined by the graph Laplacian of two high throughput proteome screenings. Identifies module regulators as singularities within the manifolds (the so-called singular manifolds). And reinterprets the characteristic IBD nonlinear dynamics as compensatory responses to perturbations on those singularities. Thus, we could control the evolution of the disease status by reconfiguring particular setups of immune system to an innocuous target state.

## Introduction

The response of a living system facing threats is a decision-making process depending on multiple factors. Some of those factors such as limited resources or energetic cost shape the different phenotypic status of a disease. Hence, the identification of “transition gates” (hereinafter referred as singularities) between those phenotypic phases opens a path to eventual therapeutic interventions with the aim of reconfiguring the system to a normal status. In particular, these status can be expressed as different grades of inflammation when the immune system of the gastrointestinal tract reacts to the presence of harmful stimuli. If the inflammatory conditions of the colon and small intestine become chronic, then, they are all generally grouped under the heading of Inflammatory bowel disease (IBD). IBD is an intestinal disease of unknown cause whose prevalence is in continuous growth at present. Crohn’s disease (CR) and ulcerative colitis (UC) are the main variants of IBD. UC, for instance, is characterised by chronic inflammation and ulceration of the lining of the major portion of the large intestine (colon). According to the European Medicines Agency UC presents a prevalence of 24.3 per 100,000 person-years in Europe. That means there are between 2.5 and 3 million people having IBD in the European Union (1) and this figure could be increased to 10 million worldwide in 10 years. The majority of patients are diagnosed early in life and the incidence continues to rise with 178,000 new cases of UC each year; therefore, the particular effect of UC on health-care systems will rise exponentially. IBD is a continuous threatening state that may produce aberrant cell proliferation leading to broad epithelial alterations such as dysplasia. This scenario of chronic active inflammation in patients with UC increases the risk for the development of colorectal carcinoma (CRC) and often requires total colectomy in case of intensive medical treatment failure or positive by high-grade dysplasia. Thus, detecting and eliminating or even reverting precursor dysplastic lesions in IBD is a practical approach to preventing the development of invasive adenocarcinoma. Currently, practitioners make decision on what new therapy to be used in such cases based only on their grade of expertise in inflammatory domains. Due to the limited and largely subjective knowledge on this pathology, we were encouraged to seek the biomarker(s) whose symbiotic actions influence the molecular pathogenesis of the risk for colorectal cancer in IBD. In this work, we develop a framework for identifying those dynamical players by abstracting the IBD progression. Since this disease can be naturally observed under the prism of a “phase transition” process (2), we sample the expression profiles of patients from a manifold with singularities; evaluating the functions of interest to the IBD status geometry near these points. In IBD, several studies have identified mucosal proteomic signatures in both active and inactive patients (11). Thus, we hypothesised that protein biomarkers could help predict the therapeutic response in patients with different status of the disease progression. Given this assumption, we construct weighted protein co-expression graphs of each variant of the disease by means of a proteomic high-throughput screening consisting of two cohort of 20 patients each (replica 1 and replica 2). Next, we localise key players of any identified modules (3) that are relevant to the IBD status, i.e., control, active and quiescent. Then, we use functions associated to the eigengenes of selected proteins across patients (3) to describe the potential of protein expressions with respect to the disease status. And we lay emphasis on the behaviour of the graph Laplacians corresponding to points at or near singularities, where different transitions of disease come together. This scenario enables the identification of potential drug targets in a protein-coexpression graph of IBD, accounts for the nonlinear dynamics inherent to IBD evolution and open the door to its eventual regression to a controlled trajectory (4–6). Overall, this manuscript envisages providing clinicians with useful molecular hypotheses of dysplastic regulation and dynamic necessaries prior to make any decision on the newest course of the treatment of individual patients in IBD. And ultimately accelerate drug discovery in health-care system.

## Results

### WGCNA Identifies Novel Immune Drivers Causing Singularities in the Status of Disease Progression

Intuitively, one might envisage the progression of a disease as a set of immune subsystems influencing each other as response to an undesired perturbation of the normal status. In this exchange, there exist specific configurations that cause the entire system changes its behaviour or collapse. We were, then, interested in identifying the potential modulators of IBD state whose interaction may explain the disease progression as a system instead of simply investigating disconnected drivers dysregulated in their expression levels. To this end, we provide protein significance measures of protein co-expression graphs of each type of IBD disease (CR and UC). Furthermore, those are simultaneously topological and biologically meaningful measures since they are defined by *cor*|*x*_*i*_, *S*|^*ξ*^ with *ξ* ≥ 1 and based on the clinical outcomes of two proteomic samples (SI Text) that capture the IBD phenotype or status, i.e., control, active and quiescent. We fixed this status as a quantitative trait defined by the vector *S* = (0, 1, −1). And applied the Weighted Gene co-Expression Network Analysis (7, 8) between the two samples, herein considered as replica 1 and replica 2 cohorts. For the sake of clarity, we only show the results obtained for the UC graph (Fig. 1). The matched inspection of its expression patterns in connectivity (Fig. 1A), hierarchical clustering of their eigengenes (9) and its eigengene adjacency heatmap (Fig. 1B-D) suggests that the most correlated expression patterns with the IBD status (Fig. 1E) are highlighted in greenyellow (97 proteins) and green (215 proteins) respectively. Whereas in CR those coloured in magenta (123 proteins) and midnightblue (41 proteins) yielded the highest correlations (Fig. S1). Nevertheless, we only kept green (UC) and magenta (CR) patterns since the others were not well preserved in the graph corresponding to the validation cohort (Fig. 2B-C and Fig. S2). Next, we wonder about the biological functions these patterns of similar protein expression to the IBD status were enrich of. To response this question, the weighted co-expression subgraph of the green (resp magenta) pattern was interrogated (Fig. 1F) using GO (10). As we expected, the green and magenta expression patterns present overabundance of IBD-related with multiple processes that are essential for the disease progression (Tables 1 and S1) such as innate immune response in mucosa, positive regulation of B cell proliferation (Fig. 3C) or inflammatory response (Fig. 3E). Complementary, pathways highly related to the disease progression such as intestinal immune network for IgA production (hsa04672) (11) or known pathways such Inflammatory bowel disease (hsa:05321) are also found (Table S2). Some surprising terms, especially other diseases such as tuberculosis, influenza A, and diabetes, appeared more enriched than IBD maybe suggesting shared signalling pathways communicating them all. In addition, the intersection of dysregulated protein sets involved in such pathways between UC and CR is very low (Fig. 3 and Fig. S3). As positive control the differential expression of well-known proteins participating in IBD such as CAMP or LYZ in UC and LCN2 or IFI16 both in UC and CR are also detected. In particular, the set composed by the proteins STAT1, AZU1, CD38 or NNMT in UC or DEFA1, IGHM, PGLYRP1 and ERAP2 in CR are robustly associated with the status of the disease (see Tables 1 and S2). These nodes, and other frequently-occurring nodes such as SYK and CD74, are attractive candidates for experimental verification. Some of these proteins work in tandem, with control sets formed by S100A9 and S100A8 identified in innate immune response process. Strikingly, some proteins such as CTSH presented in processes such as adaptive immune response or regulation of cell proliferation show similar expression between active and quiescent UC status (Fig. S4). Moreover, in CR the regulation of immune response process displayed similar expression in the active and quiescent status for all its proteins mostly belonging to the immunoglobulin heavy variable protein family that participate in the antigen recognition (Fig. S5). It should be also pointed that proteins more expressed in the quiescent than in the active status such as SYK or IGKV2-30 are mainly identified during the CR progression (Fig. S6). Upon the performance of this functional analysis, a total of 37 WGCNA candidate proteins were selected as disease-relevant within the green module of UC. Whereas the selected proteins in the case of the magenta module in CR were 19 (SI text).

**Table 1.** Analysis of the biological functions enriched in GO. We provide the proteins associated to the green module inferred by the WGCNA UC analysis along their multi-test corrected p-values.

| Biological Process in GO | Proteins | Green Module |
| --- | --- | --- |
| Immune response | HLA-DQB1, CD74, CEACAM8, SERPINB9, IGLV3-5, IGHV3-53 | $p = 1.3e^{-5}$ |
| Response to drug | LDH, NNMT, ADA, STAT1, ASSQ, DAD1, LCN2, CD38, SRP68 | $p = 2e^{-3}$ |
| Innate immune response in mucosa | S100A9, DEFA1, S100A8, SYK, IFI16 | $p = 6e^{-3}$ |
| Adaptive immune response | CTSH, TAP1 | $p = 4e^{-2}$ |
| + Regulation of cell proliferation | KRT6A, NOP2, CTSH, CAMP, RAC2, MZB1, PRTN3, NAMPT | $p = 7e^{-2}$ |
| Cell proliferation | CD74, REG1B, CDV3, ISG20 | $p = 1.6e^{-2}$ |
| Inflammatory response | AZU1, LYZ, ABCF1 | $p = 8e^{-3}$ |

**Fig. 1.**
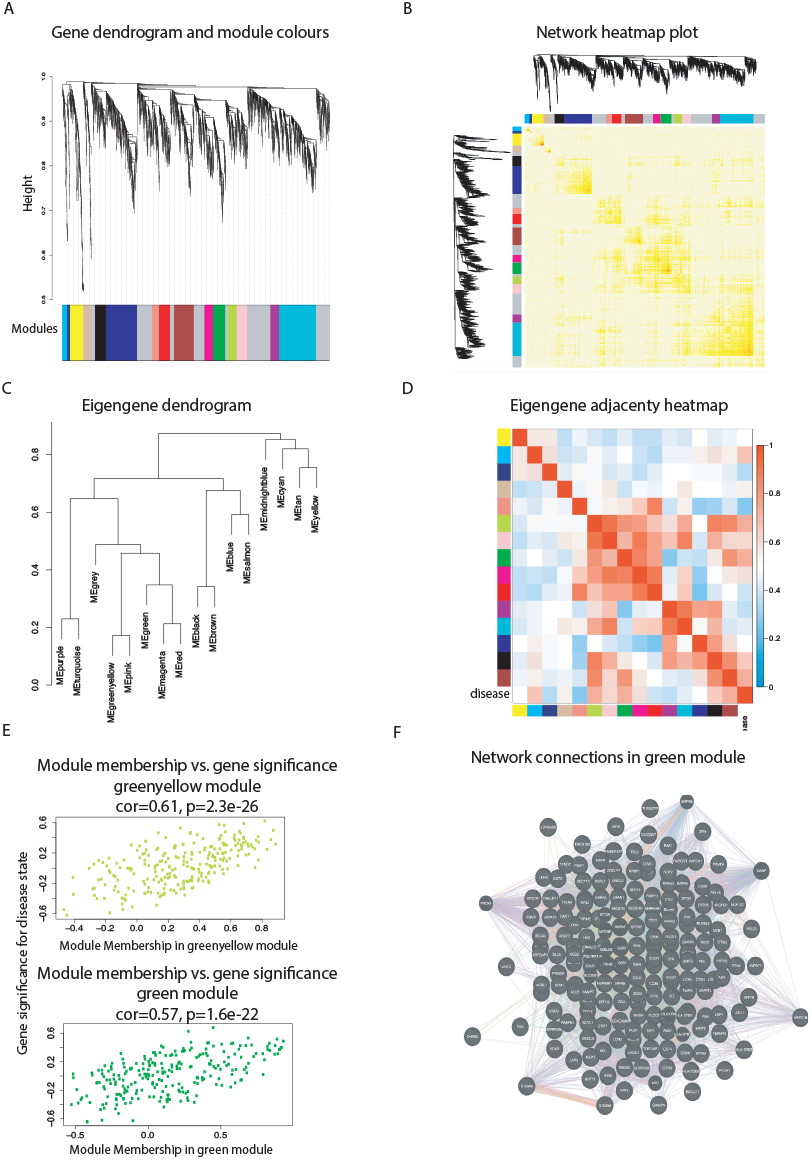
Overview of the protein coexpression network analysis in UC. (A) Hierarchical cluster tree of the 3,910 proteins analysed. The colour strips simply display a comparative overview of module assignments by means of a dynamic method to branch cuttings introduced in (12). Modules in grey are composed by “housekeeping” proteins. (B) Topological Overlap Matrix (TOM) plot (also known as connectivity plot) of the network connections. We rank the proteins in the rows and columns following the clustering tree classification. The colour scheme smoothly ranges from faint to thick nuances according to a low or a higher topological overlap. Typically data clusters along the diagonal. We also include both the cluster tree and module assignment that lie on the left and top sides respectively. (C) Hierarchical clustering dendrogram of the eigengenes calculated by the dissimilarity measure *diss*(*q*_1_, *q*_2_) = 1 − *cor*(*E*^(*q*1)^, *E*^(*q*2)^) (12). (D) Eigengene network visualisation that amounts to the relationships among the modules and the disease status. The eigengene adjacency *A*_*q*1, *q*2_ = 0.5 + 0.5*cor*(*E*^(*q*1)^, *E*^(*q*2)^ (12) (E) Protein significance versus module membership for disease status related modules. Both measurements keep a high correlation enhancing the strong correlations between the IBD progression and the respective module eigengenes (i.e. greenyellow and green). (F) Graph of the green module enriched with subgraphs functionally involved in IBD progression.

**Fig. 2.**
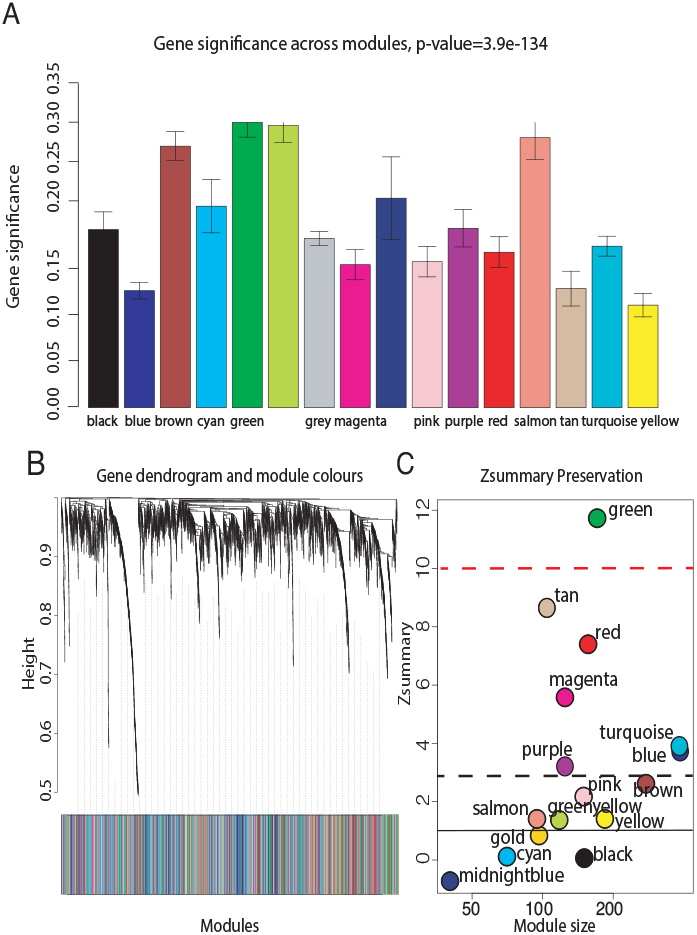
Significance and preservation of UC graph modules. (A) The module significance (average protein significance) of the modules. The underlying protein significance is defined with respect to the patient disease status. (B) The consensus dendrogram for replica 1 and replica 2 UC co-expression graphs. (C) The composite statistic *Zsummary* (Eq.9.1 in (12)). If *Zsummary* > 10 the probability the module is preserved is high (13). If *Zsummary* < 2, we can say nothing about the module preservation. In the light of the Zsummary, it is apparent there exists a high correlation with the module size. The green UC module shows high evidence of preservation in its two replica graphs.

**Fig. 3.**
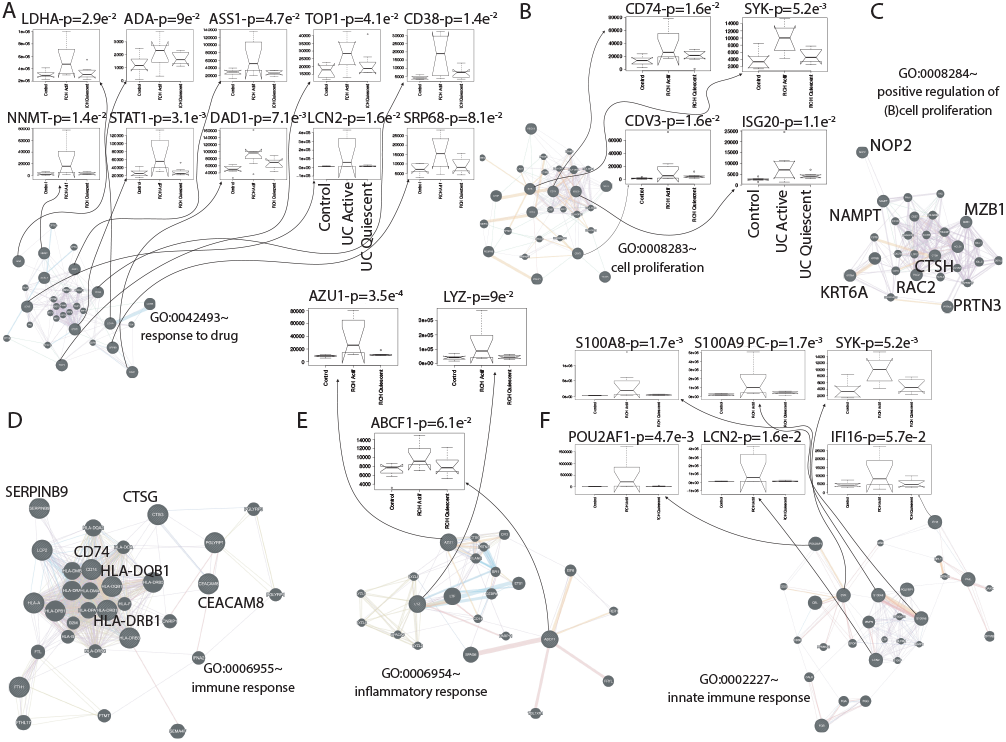
Representative repertoire of graphs by enriched functions that are well preserved in UC. (A) Response to drug. (B) Cell proliferation. (C) Positive regulation of B cell proliferation. (D) Immune response. (E) Inflammatory response. (F) Innate immune response. The protein interaction graphs were constructed using Genemania (14). Edge colouring of the graphs amount to: purple stands for Co-expression, orange for Predicted, light-blue for Pathway, light-red for Physical interactions, green for Shared protein domains and blue for Co-localisation. Boxplots of expression for the most attractive drivers respect to UC status are indicated by floating arrows at the top of each subgraph. Initial P upon the boxplot title amount to p-value associated to the IBD status correlation, whereas PC stands for positive control, i.e., proteins already described as IBD related. For the sake of simplicity, we highlighted some candidates simply by its identifiers. See Fig. S0 for more details on networks.

In the light of such computational predictions, we hypothesise the effectiveness of dynamically addressing short-term actions on the identified novel immune drivers in changing the IBD status. Remarkably, the experimental identification of such modifications would be not tractable otherwise.

### IBD Progression Can Be Geometrically Interpretable as the Intersection of Manifolds with Boundaries

Since we seek to physically control certain changes in the disease, it seems plausible to enhance the important role the geometry plays in interpreting the nature of the IBD dynamics. In many practical cases, i.e. some image-based problems, data explicitly lies on a manifold. The data are not only highdimensional, but also highly nonlinear in the most of biological systems. Particularly, that is the case of the UC and CR graphs analysed above. Nevertheless, they are endowed with a true dimension much lower than the number of features. Manifolds, herein noisy Riemannian manifolds with a measure, provide then a natural framework to understand the structure of high-dimensional data in Biology.

Notably, the data geometry of IBD status can be learnt using in an on top space the differential structure defined by those proteins selected as candidates by the WGCNA method. Each IBD status is abstracted as a Riemannian manifold, and construct the graph Laplacian based on these same candidates to figure out how the system gets cross from one to other status. Hence, we describe the domain of IBD progression as intersection of three manifolds, i.e., Ω_*c*_, Ω_*q*_ and Ω_*a*_ (Fig. S7). Let define *f*: *𝒢* → ℝ as the estimate function generated by the eigengenes associated to the WGCNA candidate proteins (Fig. S8 and S9) on the UC/CR graph *𝒢*. Next, for each couple (*i, j*) in *𝒢* we smoothly can construct a functional by:

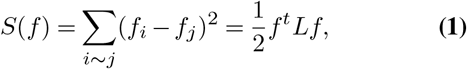

Then data derived from *𝒢* can be naturally represented on a manifold preserving adjacency by optimising the following problem:

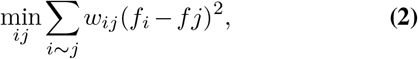

the expression 2 is scaled by the edge weight matrix of *𝒢*, 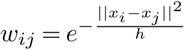 and *f*: *𝒢* → ℝ as introduced above. From (15) the associated best solution is provided by:

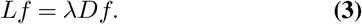

That is, the UC/CR graph *𝒢* can be represented on a manifold 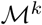 via the graph Laplacian eigenmaps. In particular, the eigenmaps of the initially mentioned eigengenes of each WGCNA candidate proteins lay on points nearby the manifold changes its structure, i.e., singularities in the manifolds intersection. Later, we inspect the behaviour of the graph Laplacian to define a potential where the dynamic of IBD status through the manifolds maybe modelled.

Let’s have a look to the construction of the graph Laplacian *L*_*n,h*_, which must be suitably scaled by 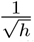. To this, we use the Gaussian kernel with bandwidth *h* (SI Text) on the 37 candidate proteins (see previous section) data selected in UC (resp. 19 in CR). Yet, we identify the cross from control to active status of the disease with an intersection-type singularity since it can be naturally considered a “phase transition” as described in *Belkin et al. (2, 15)*. Whereas the active-quiescent pass is interpreted as an edge-type singularity since the manifold sharply changes direction (i.e., from disease to control-like status). We are particularly interested in the former scenario, which involves the intersection of the two different manifolds Ω_*c*_ and Ω_*a*_. Thus, for a given point *x*_1_ ∈ Ω_*c*_ consider its projection *x*_2_ onto Ω_*a*_ and its nearest neighbour *x*_0_ in the singularity. If *n*_1_ and *n*_2_, are the directions to *x*_0_ from *x*_1_ and *x*_2_ respectively and *D*_1_ and *D*_2_ are the corresponding distances calculated as the KullbackLeibler divergence (16) between each status. From (2) *L*_*h*_*f*(*x*_1_) can be approximated by 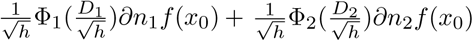. Note Φ_1_,Φ_2_ are scalar functions meeting in form and type singularity. Both functions are explicitly calculated in 4 for the intersection-type singularity. To perform this calculation, we apply Theorem 2 pg. 37 in (2) first to establishing the conditions to analyse the behaviour of the graph Laplacian near the intersection of the two 8-manifolds 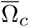 (i.e. the interior of the smooth Ω_*c*_ eventually with boundary) and 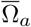 embedded in ℝ^20^. A smooth manifold of dimension *l* (≤ 7) comes out from the intersection of Ω_*c*_ ∩ Ω_*a*_. Furthermore, we fix the restricted function 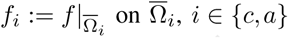 of a continuous function f defined over 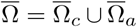 to be *C*^2^*-continuous*. And finally we consider two points, *x* Ω_*c*_ nearby the intersection and *x*_0_ being its nearest neighbour in Ω_*c*_ Ω_*a*_ and their projections *x*_1_ (resp. *x*_2_) in the tangent space of 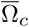 at *x*_0_ (resp. in the tangent space of 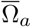 at *x*_0_). Hence, to a proper distance 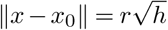 with a sufficiently small *h*, we may have

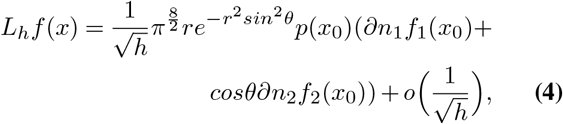

where *n*_1_ and *n*_2_ are the unit vectors in the direction of *x*_0_ − *x*_1_ and *x*_0_ − *x*_2_, respectively, and *θ* is the angle between *n*_1_ and *n*_2_ measured as the disease incidence during the cohorts recruitment (see Table 2). From equation 4, *L*_*h*_*f* (*x*) can be implicitly transform into 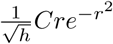 to a constant *C* depending on the derivatives of *f* and the position of *x* on or nearby the intersection (see (2, 15)).

This result naturally defines a potential (Figs. 4 and S10) properly describing the IBD control-active set, which reinforce the hypothesis exposed in the previous section. There, we claimed how therapeutic targets within the candidate derived from the proteomic coexpression dataset could be effective in the control of certain disease dynamics. The existence of this potential ultimately confirms the hypothesis by simply conducting further dynamical enquiries on it as is described in the next section.

**Table 2.**
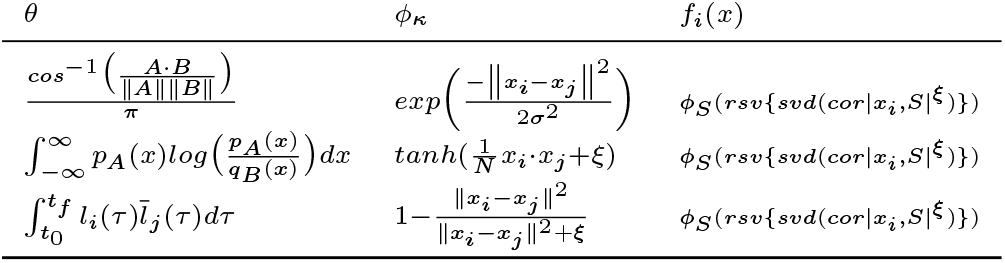
Manifolds incidence *θ*, kernel search *φ*_*κ*_ and eigengene form *f*_*i*_(*x*). First row corresponds to the optimal selection.

**Fig. 4.**
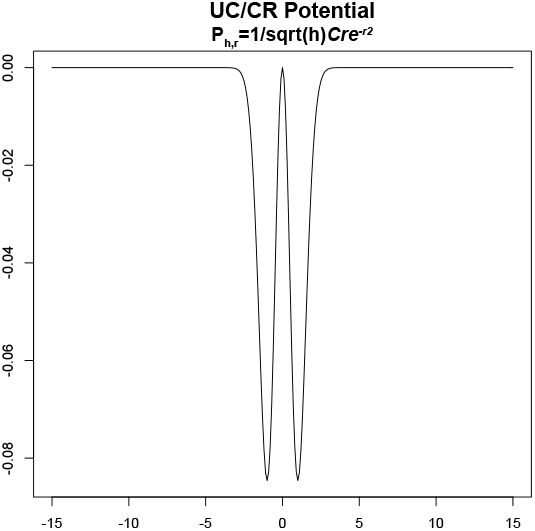
Potential of the UC control-active set. The geometry described, coupled with the abstracted disease-related dynamics through this potential, can be used to prioritise therapeutic interventions.

### Therapeutic reconfiguration of the IBD Complex Space

Finally, we want to identify temporary actions in the expression of the selected therapeutic targets that potentially control the dynamics of IBD status. That would definitely state our initial hypothesis. To this end, we followed Cornelius et al. (4) in the construction of a control perturbations in an eight-dimensional system. This type of control consists of a particle defined as the eigengene associated with one of the selected proteins across patients in their corresponding manifold of definition, i.e., Ω_*c*_, Ω_*q*_ and Ω_*a*_. We evaluate, then, its expression in function of the potential calculated in the previous section. The system of ordinary differential equations (ODEs) describing this particle at or near an intersection-type singularity as introduced above maybe simplified as:

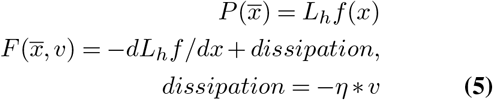

Right-hand side of ODEs defining system, according to Newton’s second law, i.e., if *y* = (*x, v*):

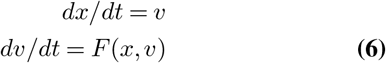

Before solving the systems of ordinary differential equations defined in 6, we optimised the dissipation parameter *η* (Fig. 5) by performing a 100 bootstrap simulations (17). If our intuition is correct, we would expect to determine the class of control perturbations that drive a particle near an undesired stable point to an innocuous target state of the disease. In effect, with *η* = 0.1, the system has two stable fixed points, one at *x* < 0 and the other at *x* > 0 (with *dx/dt* = 0 and *sd* ∼ *O*(6.5*e*^−8^)). If we continue the steady state of the system from a starting point near to the origin (i.e. 0.075), there exists a bifurcation in −0.08 that well separates the two “basin of attraction” *x*_*D*_ and *x*_*C*_ representing the bistable domain (Fig. 6) of the IBD status (SI Text). This scenario holds the non-linear dynamics implicit in the progression of IBD. Importantly, it captures the iterative interventions on the expressions of the previously selected proteins required to effectively brings the system to a non-active status from an uncontrolled path of the IBD status. Let *y*_*D*_ and *y*_*C*_ be the positions of stable fixed points minimising *L*_*h*_*f* (*x*) at *x*_*D*_ and *x*_*C*_ respectively. We first fix an initial state near *x*_*D*_ to take then a state in the basin of *y*_*D*_, and try to drive it to the basin of *y*_*C*_. This resulted in a pertinent class of control perturbations highlighted as red arrows on the left hand side of Fig. 6. Complementary, we also calibrated the class of compensatory perturbations causing the dynamic of a state on an unbounded orbit, i.e. *x* → + ∞, be driven onto the basin of *y*_*C*_. Similarly to the previous case, this class is also represented by a red arrow, but this time on the right hand side of Fig. 6. Specifically, we find that we are able to rescue the same pre-active or quiescent status above with an average distance in norm of *O*(*e*^−20^) of a feasible target status. These interventions affect a small number of proteins that in turn are multi-target, which is highly desirable provided IBD status progression is believed to be in a multi-facet cellular components synchrony. The dynamics of UC and CR share a unique pattern, but involving a few different proteins in the reconfiguration of their systems. Consequently, our initial hypothesis would be proved by programming actions performed by the IBD potential on the promising therapeutic targets within the co-expression dataset. Yet, to highlight how an experimental validation by systematic screenings of our proteomic data would be unaffordable; what enhances the potential of our methodology.

**Fig. 5.**
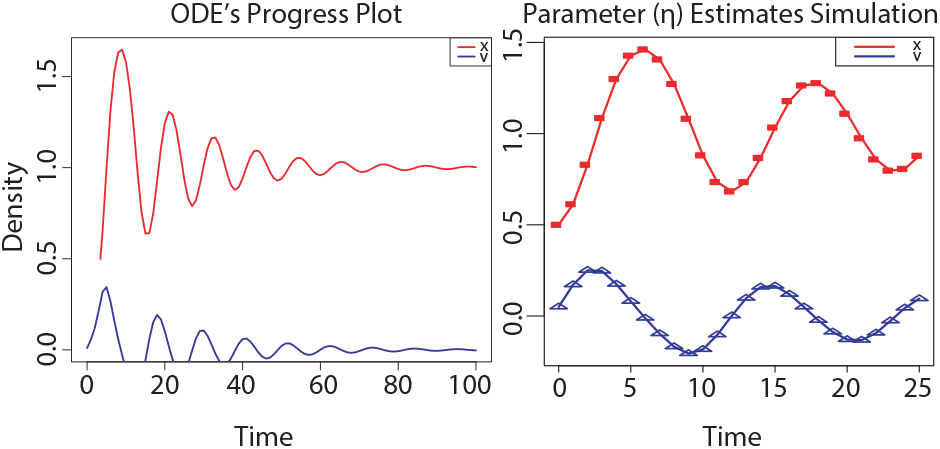
Fitting our model to data. (A) Time plot of the ODE’s system associated to IBD status. (B) Curves show the model for the best estimated parameters released upon a 100 bootstrap iterations, and the symbols depict the data.

**Fig. 6.**
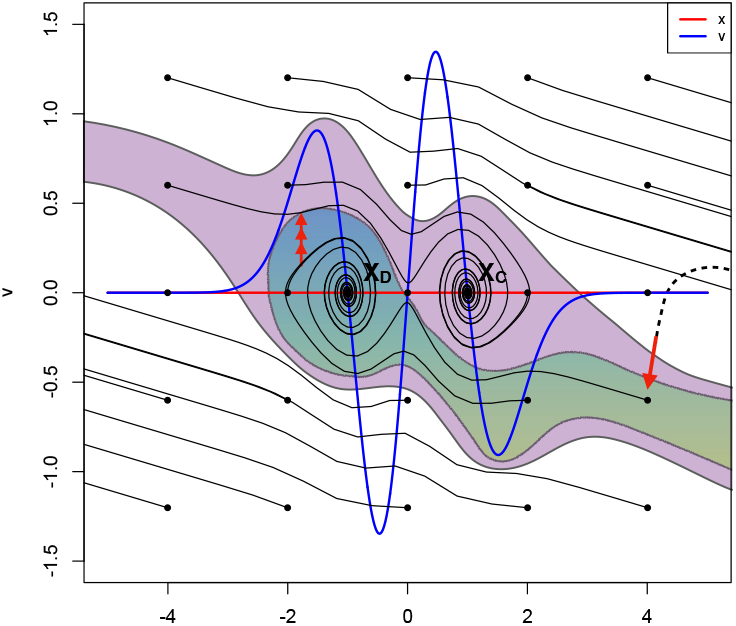
Illustration of the control process in two dimensions. The basins of attraction of the stable states *x*_*D*_ (Disease) and *x*_*C*_ (Control or Latent) are highlighted in green and violet respectively. White corresponds to unbounded orbits (left and right hand side red arrows). Iterative construction of the perturbation for an initial state in the basin of *x*_*D*_ with *x*_*C*_ as a target (Left hand side), and for an initial state on the right side of both basins with *x*_*D*_ as a target (Right hand side). Dashed and continuous lines indicate the original and controlled orbits, respectively. Red arrows indicate the full compensatory perturbations. Individual iterations of the process are shown in the insets (for clarity, not all iterations are included). Figure adopted from (4).

## Conclusions

We introduce a systematic strategy to identify potential immune drivers whose variation in expression can explain the different status displayed in the evolution of two cohorts of IBD patients. To this end, we model IBD status by the intersection of special geometric varieties called manifolds. And leverage the graph Laplacian to identify points, on or nearby their intersection, with highly specific module regulators of protein co-expression graphs. These graphs were constructed based on high-throughput proteome screenings of the two samples of IBD patients, i.e., replica 1 and replica 2 cohorts. Then to make this methodology more biological meaningful, we can test our predictions by means of experiments that study how specific interventions influence the reprogramming of IBD status, either via high-throughput sequencing surveys of validation populations (18) or by immunofluorescence specific to given antibodies (19). The comparison between theory and experiment will provide insight into the functional constraints of immune system in the recognition of bio-drivers varying when facing an intestinal chronic inflammatory threat.

The continuous trade-off amongst the available resources in living systems determines, in a certain way, their response to many situations of stress. To put this mechanism of response in motion, organisms tend to deploy a large diversity of components, such as cell types or proteins (20, 21), each sensitive to a small section of their domain. For example, the colon supports the interplay between reactive oxygen and nitrogen species overproduction or cytokines growth factors, that collectively represent a pivotal role of behaviourally aspects in IBD-induced carcinogenesis (22). Likewise, the role of the binomial composed by the innate and adaptive immune system in IBD therapies involves dozens of feedback loops invoking and sustaining chronic inflammation. However, how the immune system sparks a particular response to repel an IBD threat remains confusing, though it is thought to be immune and non-immune based, with an accepted role of the gut microbiota and non-immune derived cells of the inflammatory cascade including chemokines and inflammasomes (23). In this case, the multifaceted information process limits the repertoire of the immunological machinery. To deal with those specific environmental forces, living systems wisely prioritise their resources in accordance with their costs, and constraints (24). In this work, we have shown how the immune system response in IBD is subjected to such combination of elements, what could fix the status of IBD during its evolution. Our finding reproduces an optimal framework to detect novel immune driver-type specific to IBD status and relates it to the concept of their non-linear dynamics nearby singular topological settings (25). In this context, limiting regions of phenotypic space are modelled by means of Riemannian manifolds revealing themselves suitable to reflect the important competition between resources and costs when that eventually exceeds a vital threshold in IBD. In general this unbalanced response forces the system to drive trajectories of IBD patients to undesirable status of disease (26), our model would lead those trajectories to a region of initial conditions whose trajectories converge to a desirable status –similar to the “basin of attraction” introduced in (4). The connection between the identified immune drivers-type specific and their implication in the evolution of IBD network status becomes even clearer analysing the results yielded by our dynamical model where the pass from one to other status could explain the synergy between innate/adaptative immune resources and their energetic cost and grow in relation to their success in securing resources. Although this study is a characterisation by oversimplification of the adaptive immune system detecting dysplastic lesions in IBD, we expect that our methodology and results will be instrumental also for other diseases and thus have a more wider application for the biomedical field and associated health care systems.

## Materials and Methods

The calculations related to these sections were implemented using scripts based on R for weighted graphs analysis (wgcna package (27)), in-house Matlab^©^ (2011a, The MathWorks Inc., Natick, MA) functions for the analysis of singularities on manifolds and Python for nonlinear optimisation of control perturbations.

### Data

Samples of 30 *π*g of protein prepared from five group of patient biopsies from sigmoid colonic inflamed mucosa extracted from two cohorts (CTRL, active Crohn, quiescent Crohn, active RCH, quiescent RCH; 8 samples by group).

### Proteomic Screening

LC-MS/MS acquisition in samples of 30*µg* of 3,910 proteins was prepared from the groups of patient biopsies running on a NUPAGE 4-12% acrylamide gel (Invitrogen) and stained in Coomassie blue (Simply-blue Safestain, Invitrogen). Peptides and proteins identifications and quantification by LC-MS/MS were implemented by Thermo Scientific, version 2.1 and Matrix Science, version 5.1.

### WGCNA

We adopted the standard flow of WGCNA (7) to constructing the protein graphs of UC and CD, detecting protein modules in term of IBD status co-expression and detecting associations of modules to phenotype i.e., control, active and quiescent disease with a soft-threshold, *ξ*, determined according to the scale-free topology criterion (SI Text). Gene ontology analyses coupled with bioinformatics approaches revealed drug targets and transcriptional regulators of immune modules predicted to favourably modulate status in IBD.

### Notes on the Limit Analysis of Graph Laplacian on Singular Manifolds

That the graph Laplacian converges, in the interior points of its domain, to the Laplace-Beltrami is already known by generalisation of Fourier analysis (2, 28). Moreover, the eigenfunctions associated to this operator act as natural basis for the *L*_2_ functions used to represent data on the manifold (29). The points nearby phenotypic changes of the disease space are not interior though. Hence, we must draw upon the definition of limit from (2) if we want to analyse the behaviour of our infinite graphs Laplacian. Specifically, *L*_*n,h*_, *n* = *∞*, *L*_*h*_*f* (*x*) in UC and CR, when *x* is on or nearby a singular point, *h* is small and the function 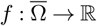, is fixed.

For a fixed *h* we define *L*_*h*_ as the limit of *L*_*n,h*_ as the amount of data tends to a very high scale:

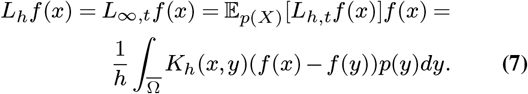

The graph Laplacian is scaled by the Gaussian kernel *K*_*h*_ with bandwidth *h*. Note that *p*(*x*) is defining a piecewise smooth probability density function on 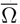. Such distribution is composed by i.i.d. 40 random samples 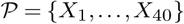 corresponding to the IBD patients. Thus, if we perform the analysis already described in Section 2 of results to the equation 7 in the intersection-type singularities on each IBD status manifold, we may deduce the equation 4.

### Nonlinear Optimisation

We learn from (4) how to optimise the interventions set needed to control the evolution of our disease model. This control procedure is iterative and consists of minimising the residual distance between the target state, *x*^∗^, and the system path *x*(*t*) at its time of closest approach, *t*_*c*_. To ensure the existence of admissible perturbations in the system herein represented by the vector expressions 9,10 and also to limit the magnitude of the solution *δx*_0_ of the optimisation problem 11, some few constraints must be introduced (SI Text). Then finding the particular solution, *δx*_0_, becomes a nonlinear programming problem (NLP) that can then be properly defined as:

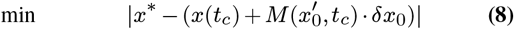

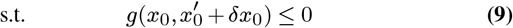

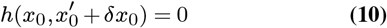

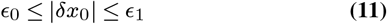

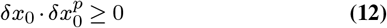

Where the matrix *M* (*x*_0_; *t*) is the solution of the variation equation *dM* = *dt* = *DF* (*x*) · *M* subject to the initial condition *M* (*x*_0_; *t*_0_) = 1. And 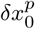 denotes the incremental perturbation from the previous iteration.

## Supporting information

Supplemental methods and results

## ACKNOWLEDGEMENTS

We acknowledge the financial support by Institut National de la Santé et de la Recherche Medicale (INSERM), Inception IBD, Inserm-Transfert, Association Franois Aupetit (AFA), Université Diderot Paris 7, and the Investissements d’Avenir programme ANR-11-IDEX-0005-02 and 10-LABX-0017, Sorbonne Paris Cité, Laboratoire d’excellence INFLAMEX.

