## Supplemental methods and results for "Singular manifolds of proteomic drivers to model the evolution of inflammatory bowel disease status"

Hatem Zaag[4], and

Eric Ogier-Denis[1]

[1]INSERM, Research Centre of Inflammation, laboratoire d'excellence Inflamex, BP 416, Paris, France

[2] Proteomic Service Jacques moneaud

[3] Inception IBD, Inc., Montreal, Canada

[4]Université Sorbonne Paris Nord, LAGA, CNRS, UMR 7539, laboratoire d'excellence Inflamex, F-93430, Villetaneuse, France

#### Full Description of Methods

Below, we describe the methods used in this work that are based on and extracted from previous results introduced by Belkin et al., 2012, Belkin et al., 2015, Horvath et al., 2008 and Cornelius et al., 2013 among others. Intellectually the work was already done independently, then, we only glued the pieces!

##### Liquid Chromatography with Mass Spectrometry (LC-MS/MS) Acquisition

Samples of 30 µg of protein prepared from five group of patient biopsies (CTRL, active Crohn, quiescent Crohn, active RCH, quiescent RCH; 8 samples by group) were run on a NUPAGE 4-12% acrylamide gel (Invitrogen) and stained in Coomassie blue (Simply-blue Safestain, Invitrogen). 3 gel plugs by sample were cut for each condition and reduced with 10mM dithiothreitol (DTT), alkylated with 55mM iodoacetamide and incubated with 20 µL of 25 mmol $L^{-1}$ $NH_4HCO_3$ containing sequencing-grade trypsin (12.5 µg $mL^{-1}$; Promega) overnight at 37°C. The resulting peptides were sequentially extracted with 30% acetonitrile, 0.1% formic acid and 70% acetonitrile, 0.1% formic acid. Peptides mixtures were pooled by sample and analyzed by an Orbitrap Fusion Tribrid coupled to a Nano-LC Proxeon 1000 equipped with an easy spray ion source (all from Thermo Scientific). Peptides separation performed by chromatography under the following conditions: Acclaim PepMap100 C18 pre-column (2 cm, 75 µm i.d., 3 µm, 100 Å), Pepmap-RSLC Proxeon C18 column (75 cm, 75 µm i.d., 2 µm, 100 Å), 300 nl/min flow rate, using a gradient rising from 95 % solvent A (water, 0.1 % formic acid) to 40 % B (80 % acetonitrile, 0.1% formic acid) in 120 minutes, followed by a column regeneration of 20 min.

Peptides were analyzed in the Orbitrap cell, in MS, at a resolution of 120,000 (at *m/z* 200), with a mass range of *m/z* 350-1550 and an AGC target of 2 x$10^5$. Fragments were obtained by high collision-induced dissociation (HCD) activation with a collisional energy of 30%. MS/MS data were gathered from the ion trap in the top-speed mode, with a total cycle of 3 seconds, with an AGC target of 5x$10^4$, a dynamic exclusion of 50 seconds and exclusion duration of 60 seconds. The maximum ion accumulation times were set to 100 ms for MS acquisition and 30 ms for MS/MS acquisition in parallelization mode.

#### Peptides and proteins identifications and quantification by LC-MS/MS

Identification procedure: The Proteome Discoverer software (Thermo Scientific, version 2.1) and with the Mascot search engine (Matrix Science, version 5.1) were used in the MS and MS/MS data processing. For the precursor ion we set to 7 ppm the mass tolerance maximum threshold and 0.5 Da for fragments. The following variable modifications (2 maximum per peptide) were allowed: carbamidomethylation (Cys), oxidation (Met), phosphorylation (Ser, Thr, Tyr), acetylation (N-term of protein). More than two missed cleavages were not considered for the trypsin protease. The MS/MS identification step performed on the SwissProt database (02/16 version) using the *Homo sapiens* taxonomy as base. Peptide Identifications were validated using a 1% FDR (False Discovery Rate) threshold calculated with the Percolator algorithm. The relative quantification of protein abundances is measured by means of Progenesis QI - Proteomics software (version 4.0, Waters) - using co-detection in order to eliminate missing values. The relative quantitation of proteins according to the five groups (CTRL, active

Crohn, quiescent Crohn, active RCH, quiescent RCH) was performed using a between subject analysis and a Hi-3 method (for which the three most abundant peptides were used for protein quantification). Abundance variations of proteins with an ANOVA p-value under 0.05 and with at least two identified peptides were further considered.

#### How to perform the Weighted Gene co-Expression Network Analysis

The WGCNA R package enables the construction of co-expression network using the normalised data of the 40 microarray-measured in the two replica samples just described in the previous section. We set the threshold power to 10 since it was the supremum in the scale-free $R^2$ fit of 0.8. Briefly, the network was created calculating topologic overlap (TO) with bicor correlation function, then genes were hierarchically clustered using 1-TO (dissTOM) as the distance measure. Initial module assignments were determined by using dynamic tree-cutting, using default parameters except deepSplit =0 , mergeCutHeight=0.20, minModulesize = 30, pamStage = FALSE, and pamRespectsDendro = TRUE, but genes were allowed to be reassigned to modules with better correlation if $p < 0.05$ for those correlations. The network type was signed, so that anticorrelated genes were not assigned to the same module. Resulting 16 (resp. 18 for CD) modules or groups of co-expressed genes for UC ranging in size from 540 genes (resp. 305 for CR) (turquoise) to 73 genes (cyan, resp. 81 in tan for CR) were used to calculate the eigengenes (MEs; or the 1st principal component of the module). MEs were correlated with the status of IBD in terms of biological trait.

In the validation WGCNA, scale free topology was achieved with beta (power) set at 17.5, and other similar to above function parameters were deepSplit = 4, minModulesize = 3, mergeCutHeight = 0.12, pamStage = TRUE, pamRespectsDentro = TRUE, reassignThresh = 0.05. The WGCNA::modulePreservation() function was used to estimate the preservation rate stats of each module between the discovery and validation cohorts of IBD patients with parameters nPermutations=30, maxGoldModuleSize=100, maxModuleSize=400. Functional analysis of over-represented processes was performed with an in-house tailored script testing for hypergeometric overlap of gene symbol membership in modules across the derivation and validation networks using the fisher.test() function achieveing to a consesus with the resulting scores obtained by means of DAVID, GORILLA, STRING, KEGG and PANTHER data bases.

WGCNA network gene-level kME values (gene expression correlation to the module eigengene for the module to which each gene belonged) were used to rule out the measured gene list to genes with kME > 0.65 and membership in any of 4 modules (resp. 2 in CR) of particular interest in UC. R code is provided online via synapse.org (https://www.synapse.org/XXXXXX).

#### Manifold Assumption of Data Representation in IBD

In many cases, i.e. some image-based issues, data explicitly lies on a manifold. The data are not only high-dimensional, but also highly nonlinear in the most of biological systems. Nevertheless, they are endowed with a true dimension much lower than the number of features. Manifolds, herein noisy Riemannian manifolds with a measure, provide a natural framework to understand the structure of high-dimensional data in Biology.

We enquire our data about how their shape might affect the notion we have of similarity. By *assuming that functions of interest are smooth with respect to the underlying geometry* the structure of real data is naturally unveiled.

Given the probability distribution $P$ on $X \times Y$ and the map $X \underset{P}{\rightarrow} Y$. We consider the function $f: X \rightarrow \mathbb{R}$ that potentially learns the shape of a data set thanks to the assumption of a penalty at $x \in X$ introduced as follows:

$$\frac{1}{\delta^k} \int_{small\ \delta} (f(x) - f(x+\delta))^2\, p(x) d\delta \approx \parallel \nabla f \parallel^2 p(x),$$

Where Laplace operator $\Delta f = -\sum_{i=1}^{k} \frac{\partial^2 f}{\partial x_i^2}$ helps to identify the total penalty by:

$$\int_X \parallel \nabla f \parallel^2 p(x) = \langle f, \Delta_p f \rangle_X.$$

Then by the manifold assumption stated in Belkin et al., 2012, *we claim* $\langle f, \Delta_p f \rangle_X$ *is small.* So then, data shape can be closely traced using such function *f*.

#### Laplace-Beltrami Operator Basis

The Laplace operator is the only differential operator invariant under translations and rotations and in a certain way acts as precursor of the basis required for the function used in the geometry of data. By operating on the circle in two dimensions we obtain similar periodic boundary conditions to $\mathbb{R}$. This result of the Fourier analysis can be generalised to infer the Laplace – Beltrami operator. But first, let's define $f: \mathcal{M}^m \rightarrow \mathbb{R}$ and $exp_p: T_p\mathcal{M}^m \rightarrow \mathcal{M}^m$ as the application to be estimate defined from the manifold of dimension $m$ *to* $\mathbb{R}$ (as describe in the previous section) and the corresponding application defined from its tangent space respectively. Then, the operator is defined as follows:

$$\Delta_{\mathcal{M}} f(p) = -\sum_i \frac{\partial^2 f(exp_p(x))}{\partial x_i^2}$$

The Laplace-Beltrami operator *provides a basis* for $L_2$functions on the manifold ordered by smoothness according to the eigenvalue.

The span of a few bottom eigenvectors $(e_1 \dots e_m)$ is a natural space of predictors for fitting data.

For example in 2D, given some protein candidates $(x_i, y_i)$ these eigenvectors enable predicting them by learning:

$$\min_{a_i, i=1\dots m} \sum_{j=1}^{n} (\sum_{i=1}^{m} a_i e_i - y_i)^2$$

**Data Representation: Graph Laplacian Eigenmaps**

Consider a Graph $G$ and its set **f** of sub functions $f_i$ defined as indicated in the previous section and this for each $(i,j) couple\ of\ nodes\ in\ the\ graph.$ We smoothly can construct a *functional* by:

$$S(f) = \sum_{i \sim j} (f_i - f_j)^2 = \frac{1}{2} f^t L f$$

Then *data derived from graph can be naturally represented on a manifold* preserving adjacency by optimising the following problem:

$$\min_{ij} \sum_{i \sim j} w_{ij} (f_i - f_j)^2,$$

the latest expression is scaled by the edge weight matrix of $G$, $w_{ij} = e^{-\frac{\|x_i - x_j\|^2}{h}}$ and $f: G \to \mathbb{R}$ as introduce above.

The associated best solution is provided by [3 Belkin Nigoyi 01]:

$$Lf = \lambda Df.$$

Therefore, data derived from the weighted gene co-expression network analysis of IBD can be smoothly represented on a manifold via the graph Laplacian eigenmaps. *But how could this be done attending to the disease status?*

#### Singular Manifolds of IBD Status

Similarly to Belkin et al. (2012), now we wondered if our system could describe a general model for a singular IBD space based on its status where to learn stratified spaces by means of some submanifolds (control, active and quiescent strata) put together as introduced in Aanjaneya et al. (2011); Bendich et al. (2012); Haro et al. (2008). In that way, we would be able to reconstruct topological invariants characterising IBD status on manifolds (see, e.g, Chazal et al. (2009); Chazal and Oudot (2008); Niyogi et al. (2008, 2011)).

The geometric problem may be briefly described as follows:

Let $\bar{\Omega}$ be the set of *three* Riemannian submanifolds, smoothly selected and being compact, of intrinsic dimension (*the different dimensionality issue may be reduced to the former case by normalizing the probability density*) 8 embedded in $\mathbb{R}^{40}$. The inner point $x \in \bar{\Omega}$, is defined as *regular* if $x$ lies strictly within *one smooth* component $\overline{\Omega_i}$ *of* $\Omega$ *with* the so-called submanifold $\Omega_i$. In other words, if $x \in \Omega_i, x \notin \Omega_j, j \neq i$ *where* $i, j \in \{control, active, quiescent\}$. Otherwise, $x$ is defined as *singular point*. We extend this notation by denominating the boundary of $\Omega_i$ as $\partial\Omega_i$. Ultimately, to make sure $\partial\Omega_i$ is smooth we need to assume $\mathcal{M}$ as a $C^2$-manifold. Now, we define $\Omega$ as the union of $\Omega_i$, and $\partial\Omega$ as the union of $\partial\Omega_i$. Furthermore, $\Omega_i$ and $\Omega_j$ may not be disjoint, then we leverage the protein candidates selected above across phenotypic changes associated to the two cohorts of IBD patients to join together the corresponding submanifold these candidates lie on with other submanifold of a different status. They intersect in their interior ($i.e, \Omega_i \cap \Omega_j \neq \emptyset$) or they are sticked along their boundaries. Among the space of possible singularities, we are basically interested in *two general classes*:

1. **intersection-type (control-active scenario):** That amounts to the in pairwise smooth intersection of submanifolds. The simplest context occurs as such an intersection is done between the interior of the submanifolds, namely: $x \in \Omega_i \cap \Omega_j, i \neq j$;

2. **edge-type (active-quiescent scenario):** Extreme distance points that lie on within the boundaries of two smooth submanifolds. That is $x \in \partial\Omega_i \cap \partial\Omega_j, i \neq j$.

In that way, we ensure the singular points smoothly knock a local manifold of lower dimension. Furthermore, the global intersection of these two intersections operates as local on the intersection of their tangent spaces. Thus, *the* ***singular manifold of the IBD status*** can be defined as the union of 3 smooth manifolds $\bar{\Omega}$ endowed with these types of singularities.

Now, let $f : \bar{\Omega} \to \mathbb{R}$ be a *piecewise-smooth function to be reconstructed from the weights of the IBD graph G*. We also need to fix some restrictions of that function to the submanifold $\bar{\Omega}_i$ such that $f_i := f|_{\bar{\Omega}_i}, i = 1, \ldots, m$. Finally to make sense of the Graph Laplacian on the interior points of every IBD status, each one of the $f_i$ must be $C^2$-continuous on them. Hence, we can smoothly construct the probability density function $p(x)$ in piecewise on $\bar{\Omega}$ along the $0 < a \leq p_i(x) \leq b < \infty$ constraint.

#### Graph Laplacian of IBD Status

Let $X = \{X_1, \cdots, X_{40}\}$ be the set of i.i.d. random samples derived from the two cohorts of IBD patients distribution with density $p(x)$ on $\bar{\Omega}$. Then, $X$ enables to build a weighted graph $G(V, E)$ of each cohort by simply matching the $ith$-sample point $X_i$ with the $ith$-vertex $v_i$ and making correspond a weight $w_{ij}$ to edge $e_{ij}$. We consider in this study the Gaussian weight function with the Euclidean distance as indicates:

$$w_{40,h}(X_i, X_j) = \frac{1}{20h} K_h(X_i, X_j) = \frac{1}{20} \frac{1}{h^{8/2+1}} e^{-\frac{\|x_i - x_j\|^2_{\mathbb{R}^{20}}}{h}}.$$

Belkin et al., 2012 suggests normalization by $\frac{1}{20}\frac{1}{h^{8/2+1}}$ in order to hold the limit analysis. With the best fitted Gaussian bandwidth of the selected protein candidates as $h = 0.0384$.

Let's define now the edge weight matrix of the graph $G$ as $W_{n,h}$ with $W_{n,h}(i,j) = w_{n,h}(X_i, X_j)$, and $D_{n,h}$ as the diagonal matrix defined by $D_{n,h}(i,i) = \sum_j w_{n,h}(X_i, X_j)$. We may want to introduce the graph Laplacian without normalization scale as the $n \times n$ matrix $L_{n,h} = D_{n,h} - W_{n,h}$. Under all these considerations we can apply the graph Laplacian to any couple of fixed smooth function $f(x)$ and point $x \in \bar{\Omega}$ as follows

$$L_{n,h}\mathrm{f(x)} = \frac{1}{nh}\sum_{j=1}^{n} K_h(x, X_j)[f(x) - f(X_j)].$$

*This is the graph Laplacian in the interior points of IBD status manifolds. But more importantly, what would its behaviour be nearby our protein candidates where a phenotypic change produces an intersection-type singularity?*

**Notes on the Limit of Graph Laplacian defined upon Singular Manifolds**

The limit behaviour of graph Laplacians should be observed under a double prism composed by the sample size $n$ and the (Gaussian) kernel bandwidth $t$. To this end, we first address the limit $h \to 0$ for data at very great scale and later gaining insights onto finite sample and rates as $n \to \infty$ by means of concentration inequalities.

In particular, herein we study how the infinite graph Laplacian $L_h f(x)$acts when $x$ lies on or nearby a singular point with an $h$ small and a fixed function $f : \bar{\Omega} \to \mathbb{R}$.

Given $h$, we introduce $L_h$ as the limit of $L_{n,h}$ as the scale of data tends to be of very high magnitude then we have:

$$L_h f(x) = L_{\infty,h}\, f(x) = \mathbb{E}_{p(x)}\left[L_{n,h} f(x)\right] f(x) =$$

$$= \frac{1}{h} \int_{\bar{\Omega}} K_h(x, y)\big(f(x) - f(y)\big)p(y)dy.$$

To best study this integral for a small $h$, we trace a local projection from the manifold to its tangent space $\pi_x: \bar{\Omega} \to T_x$. Notice that for the finite case the convergence rate for a small $\varepsilon$ is $\leq \mathcal{O}\left(2 * 24 * exp\left(-\frac{24h^{8/2+2}\varepsilon^2}{2C_v+2C_m\varepsilon h/3}\right)\right)$ [Belkin th5], where $C_v$ *and* $C_m$ are manifold dependent constants.

**ODE System for IBD Status Dynamic**

The dynamics of a complex network can often be represented by a set of coupled ordinary differential equations. We thus consider an $N$-node network whose $n$-dimensional dynamical state x is governed by

$$\frac{dx}{dt} = F(x)\,.$$

In general, the heterogeneity in the physiopathology of IBD progression tends to perturb any of its graphical representation. Let's get this modification underway at a time prior to a given $reference\ t_0$, driving it to a status $x_0 = x(t_0)$ in the attraction basin $\mathcal{B}(x_u)$ of an undesirable phenotype $x_u$. We seek then to compensate this effect by wisely identifying a perturbation $x_0 \rightarrow x_0'$ to be implemented at time $t_0$, where $x'_0$ lies on the basin of attraction $\mathcal{B}(x^*)$ of a desirable status $x^*$. For clarity, we assume that $x_u\ and\ x^*$ are fixed points, although the same holds to other types of attractors.

In the absence of any constraints there always exists a perturbation upon $x_0$ such that $x_0' \equiv x^*$. However, and as usually occurs with the real complex graphs, we need for constrained compensatory perturbations. These constraints encode biophysical conditions of the system and often invoke no modification to certain nodes, while restricting the direction and the changes outreach in others. The latter penalises the ease of removing a node versus the global assortativity of the system. We thus assume that the constraints on the eligible perturbations can be represented, as in Cornelius et al., 2013, by the vectorial expressions

$$g(x_0, x_0') \leq 0\ and\ h(x_0, x_0') = 0,$$

where both the equality and the inequality are assumed to be applied to each component.

In particular, $x_{0j}' \leq x_{0j}$ denotes constraints for accessible nodes, and $x_{0j}' = x_{0j}$ for the other nodes. Notice that non-linear constraints are also permitted in the model.

#### Optimisation of the Dissipation Parameter

We fit the dissipation parameters of our IBD model to data (i.e. $\eta$). In this analysis we randomize the initial condition, s, and parameters, p, using a normal distribution before we fit the model to the data (see Fig. 4B).

Then we calculate the estimated parameters, confidence ranges, and correlations between the parameters. Additionally, the initial condition is read from the data (instead of estimating it).

We also fit simultaneously several data sets by provide a list of data sets to the first data option. When fitting several data sets, some of the parameters could be the same across all data sets, whereas others could differ, and have a unique value in each data set (see Fig. 4B). Finally, we show how one can bootstrap the data by sampling (with recruitment) from every individual data set, and re-fit the samples using the best parameters as an initial guess (Fig. 4A). These calculations were implemented by tailored in-house functions and the R function derived from C-code GRIND [de Boer, R, 2017].

#### Compensatory Perturbations (Nonlinear Optimisation)

We seek in such spaces how iteratively compensate unwanted trajectories starting from small perturbations, as shown in Fig. 5. Given a dynamical system as introduced above and an initial state $x_0$ at time $t_0$, a small perturbation $\delta x_0$ evolves to $\delta x(t) = M(x_0, t_c) \cdot \delta x_0$ at time $t$. The matrix $M(x_0, t)$ is the solution of the variational equation ${dM}/{dt} = DF(x) \cdot M$ subject to the initial condition $M(x_0, 0) = 1$, that is normally invertible [19]. Fixed the time point $t_c$ corresponding to the perturbed orbit straight next path attaining the target $x^*$, we should be able to apply the inverse transformation,

$$\delta x_0 = M^{-1}(x_0, t_c) \cdot \delta x(t_c),$$

to identify the perturbation $\delta x_0$ in accordance with the system initial condition $x_0$ that, albeit all the possible perturbations verifying $|\partial x_0| \leq \varepsilon_1$, will approximate $x(t_c) + \delta x(t_c)$ to $x^*$ the most (Fig. 5, left hand side red arrows). This results in a good estimation for small $\varepsilon_1$, and consequently for small fluctuations in the initial conditions of the given system. Additionally, the yielded error can in fact be calculated from the matrix of second derivatives and verifed numerically by direct integration. Large perturbations can then be built up by iterating the process: every time $\partial x_0$ is calculated, the current initial state, $x_0'$, is updated to $x_0' + \partial x_0$, and a new $\partial x_0$ is calculated starting from the new initial condition (Fig. 5, right hand side red arrows).

The problem of identifying a perturbation $\partial x_0$ that incrementally moves the orbit toward the target under the given constraints is then cast as a constrained optimization problem (see Methods). Once found, the optimal incremental perturbation is applied to the current initial condition, $x_0' \to x_0' + \partial x_0$, giving the initial state for the next iteration. At this point, we test whether the new state lies in the target's basin of attraction by integrating the system $\frac{dx}{dt} = F(x)$ over a prolonged time. The procedure ends as the orbit reaches a small ball of radius $\varsigma$ around the target. Otherwise, we identify the closest approach point in a time window $t_0 \leq t \leq t_0 + T$, where $T$ is a time limit determined beforehand, and repeat the procedure. The method may be not convergent to a permissible compensatory perturbation, e.g., if the feasible region $\mathfrak{R}_f$ does not intersect the target basin $\mathcal{B}(x^*)$. Faced to these conditions, the enquiry is truncated if the method does not control the system in a pre-fixed number of iterations.

#### Supplementary Results

### Table S1. KEGG

| Biological Functions in CR | Counts | Magenta |
|---|---|---|
| Regulation of immune response | 18 | $p=2.7e^{-12}$ |
| Immune response | 24 | $p=3.7e^{-11}$ |
| Innate immune response | 23 | $p=1.4e^{-9}$ |
| Leukocyte migration involved in inflammatory response | 11 | $p=1.5e^{-4}$ |
| Antigen processing and presentation of peptide antigen via MHC class I | 7 | $p= 6.8e^{-4}$ |
| Inflammatory response | 11 | $p= 1.3e^{-3}$ |
| Immunoglobulin mediated immune response | 3 | $p= 2.2e^{-2}$ |
| Adaptive immune response | 5 | $p= 2.4e^{-2}$ |
| Negative regulation of inflammatory response | 4 | $p= 4.8e^{-2}$ |
| Innate immune response in mucosa | 3 | $p= 7.0e^{-2}$ |
| Neutrophil activation involved in immune response | 2 | $p= 9.6e^{-2}$ |

### Table S2.

| Pathways Participating in UC | Counts | Green |
|---|---|---|
| hsa04141:Protein processing in endoplasmic reticulum | 19 | $p=2.5e^{-7}$ |
| hsa03060:Protein export | 8 | $p=1.2e^{-5}$ |
| hsa05152:Tuberculosis | 10 | $p=3.9e^{-3}$ |
| hsa05164:Influenza A | 10 | $p=4.3e^{-3}$ |
| hsa04380:Osteoclast differentiation | 9 | $p=1.0e^{-2}$ |
| hsa04940:Type I diabetes mellitus | 6 | $p=2.6e^{-2}$ |
| hsa05321:Inflammatory bowel disease (IBD) | 4 | $p=2.8e^{-2}$ |
| hsa05416:Viral myocarditis | 4 | $p=2.8e^{-2}$ |
| hsa05212:Pancreatic cancer | 4 | $p=2.8e^{-2}$ |
| hsa00220:Arginine biosynthesis | 5 | $p=3.1e^{-2}$ |
| hsa05145:Toxoplasmosis | 5 | $p=3.1e^{-2}$ |
| hsa00520:Amino sugar and nucleotide sugar metabolism | 4 | $p=3.4e^{-2}$ |
| hsa04672:Intestinal immune network for IgA production | 4 | $p=4.8e^{-2}$ |
| hsa05168:Herpes simplex infection | 5 | $p=6.3e^{-2}$ |
| hsa04612:Antigen processing and presentation | 4 | $p=7.2e^{-2}$ |
| hsa05332:Graft-versus-host disease | 3 | $p=8.4e^{-2}$ |
| hsa05320:Autoimmune thyroid disease | 6 | $p=9.1e^{-2}$ |
| hsa05330:Allograft rejection | 5 | $p=9.8e^{-2}$ |
| hsa05140:Leishmaniasis | 3 | $p=9.9e^{-2}$ |

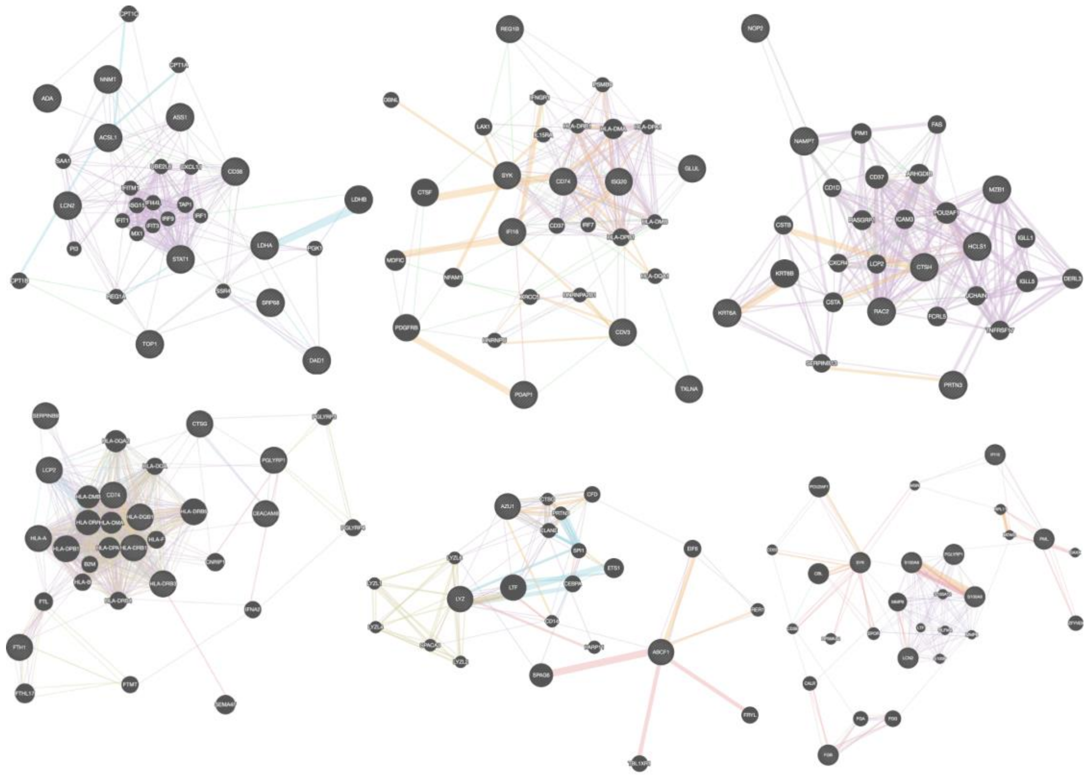

**Figure S0. View on detail of the representative repertoire of enriched graphs by functions that are well preserved in UC**. **(A)** Response to drug. **(B)** Cell proliferation. **(C)** Positive regulation of (B)cell proliferation. **(D)** Immune response. **(E)** Inflammatory response. **(F)** Innate immune response. The protein interaction graphs were constructed using Genemania (Warde-Farley et al., 2010). Edge colouring of the graphs amount to: purple stands for Co-expression, orange for Predicted, light-blue for Pathway, light-red for Physical interactions, green for Shared protein domains and blue for Co-localisation.

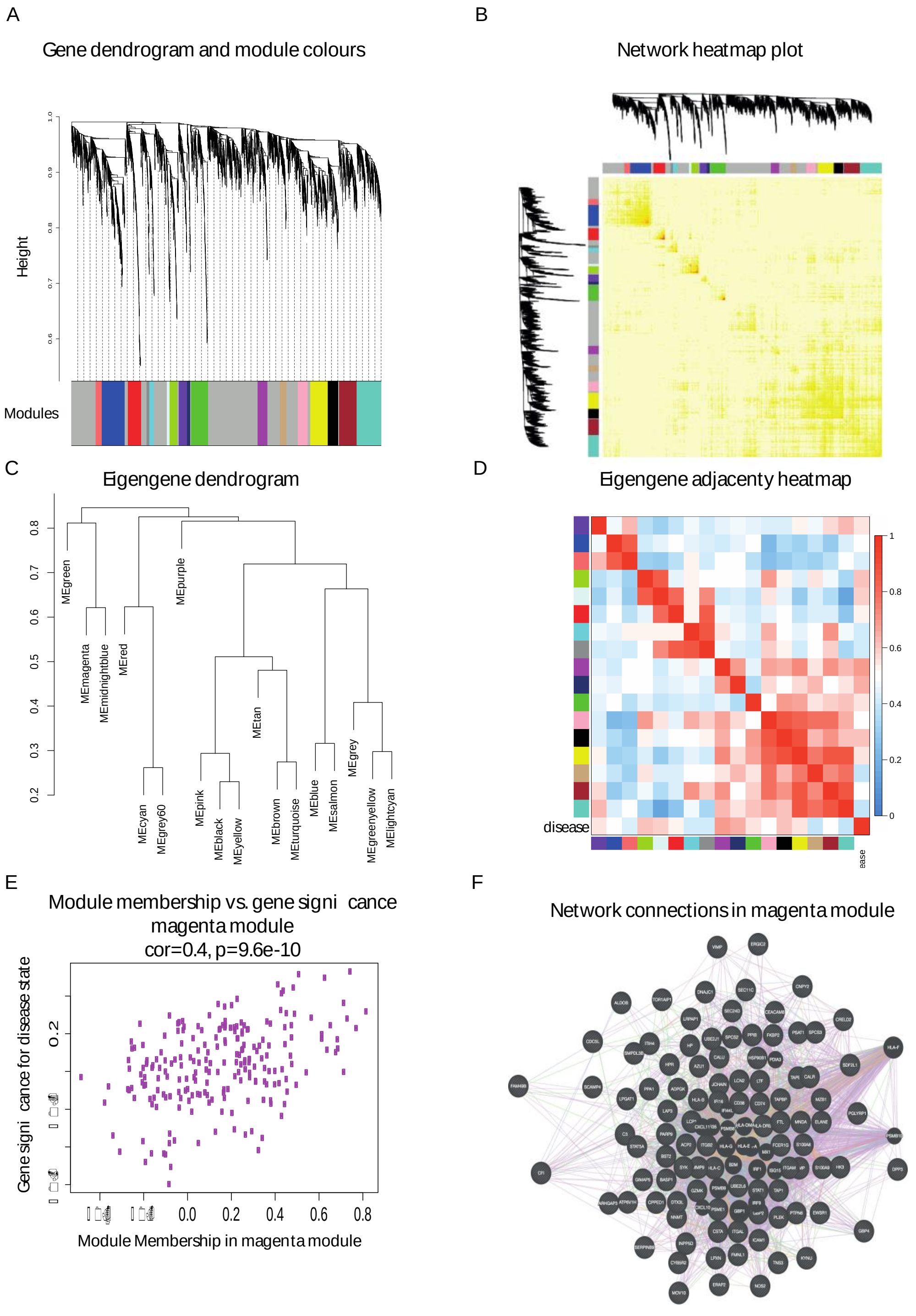

**Figure S1. Overview of the protein coexpression network analysis in CR. (A)** Hierarchical cluster tree of the 3,910 proteins analysed. The colour strips simply display a comparative overview of module assignments by means of a dynamic method to branch cuttings introduced in (12). Modules in grey are composed by "housekeeping" proteins. **(B)** Topological Overlap Matrix (TOM) plot (also known as connectivity plot) of the network connections. We rank the proteins in the rows and columns following the clustering tree classification. The colour scheme smoothly ranges from faint to thick nuances according to a low or a higher topological overlap. Typically data clusters along the diagonal. We also include both the cluster tree and module assignment that lie on the left and top sides respectively. **(C)** Hierarchical clustering dendrogram of the eigengenes calculated by the dissimilarity measure $diss(q_1,q_2) = 1 - cor(E(q_1),E(q_2))$ (Horvath, 2011). **(D)** Eigengene network visualisation that amounts to the relationships among the modules and the disease status. The eigengene adjacency $A_{q_1,q_2} = 0.5+0.5cor(E(q_1),E(q_2))$ (Horvath, 2011). **(E)** Protein significance versus module membership for disease status related modules. Both measurements keep a high correlation enhancing the strong correlations between the IBD progression and the respective module eigengenes (i.e. greenyellow and green). **(F)** Graph of the green module enriched with subgraphs functionally involved in IBD progression.

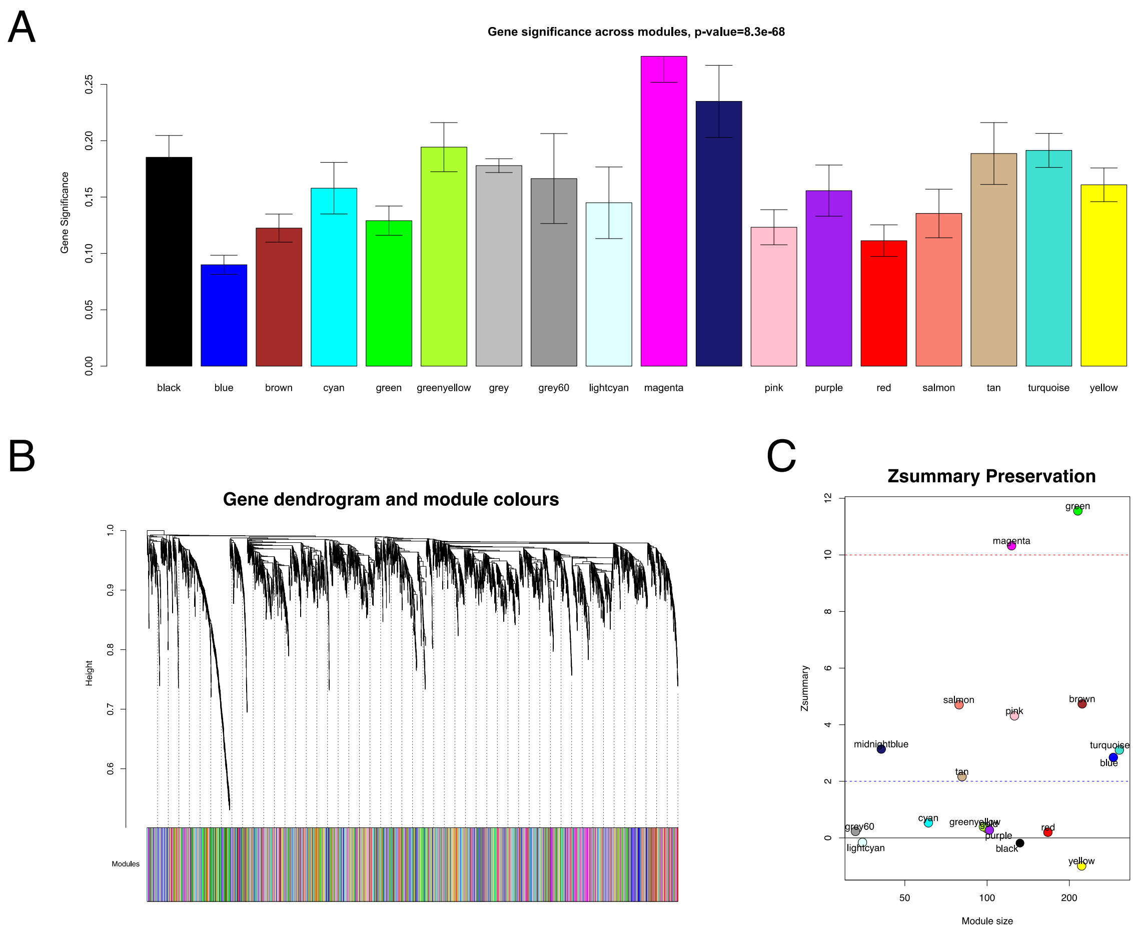

**Figure S2. Significance and preservation of CR graph modules. (A)** The module significance (average protein significance) of the modules. The underlying protein significance is defined with respect to the patient disease status. **(B)** The consensus dendrogram for replica 1 and replica 2 CR co-expression graphs. **(C)** The composite statistic *Zsummary* (Eq.9.1 in (12)). If *Zsummary* > 10 the probability the module is preserved is high (13). If *Zsummary* < 2, we can say nothing about the module preservation. In the light of the Zsummary, it is apparent there exists a high correlation with the module size. The green UC module shows high evidence of preservation in its two replica graphs.

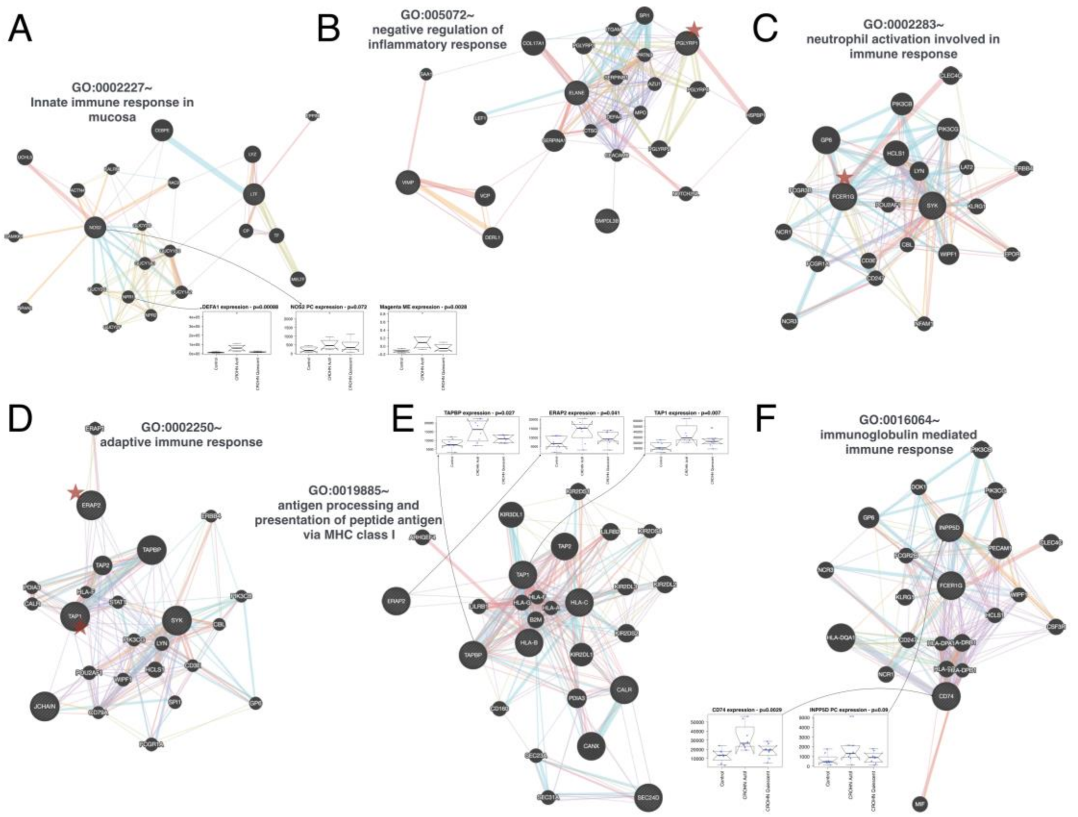

**Figure S3. Representative repertoire of enriched graphs by functions that are well preserved in CR**. **(A)** Innate immune response in mucosa. **(B)** Negative regulation of inflammatory response. **(C)** Neutrophil activation involved in immune response. **(D)** Adaptive immune response. **(E)** Antigen processing and presentation of peptide antigen via MHC class I. **(F)** Immunoglobulin mediated immune response. The protein interaction graphs were constructed using Genemania (Warde-Farley et al., 2010). Edge colouring of the graphs amount to: purple stands for Co-expression, orange for Predicted, light-blue for Pathway, light-red for Physical interactions, green for Shared protein domains and blue for Co-localisation. Boxplots of expression for the most attractive drivers respect to UC status are indicated by floating arrows at the top of each subgraph. Initial P upon the boxplot title amount to p-value associated to the IBD status correlation, whereas PC stands for positive control, i.e., proteins already described as IBD related. For the sake of simplicity, we highlighted some candidates simply by its identifiers.

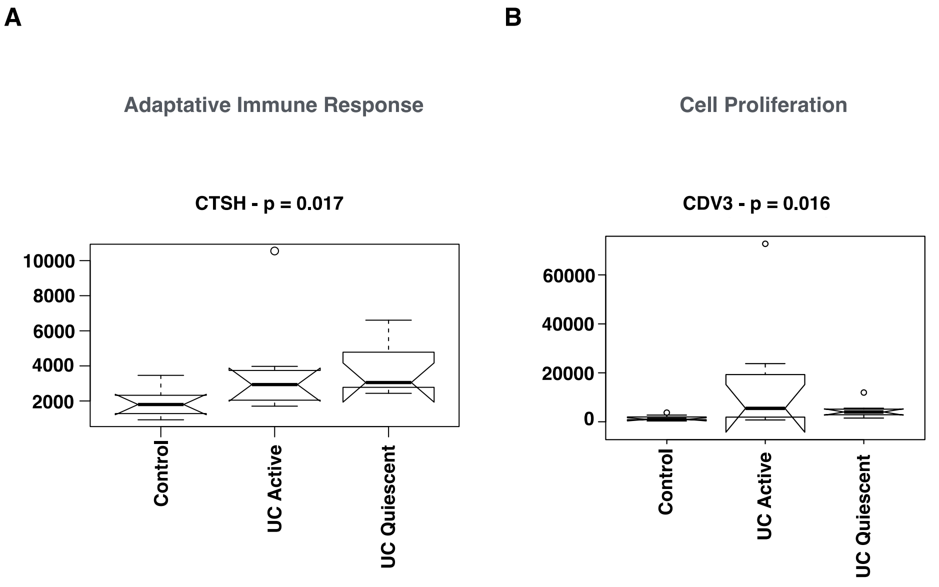

**Figure S4. Similar patterns of protein expression in the UC green WGCNA selected module.** CTSH and CDV3 barely present the same expression levels between active and quiescent UC status.

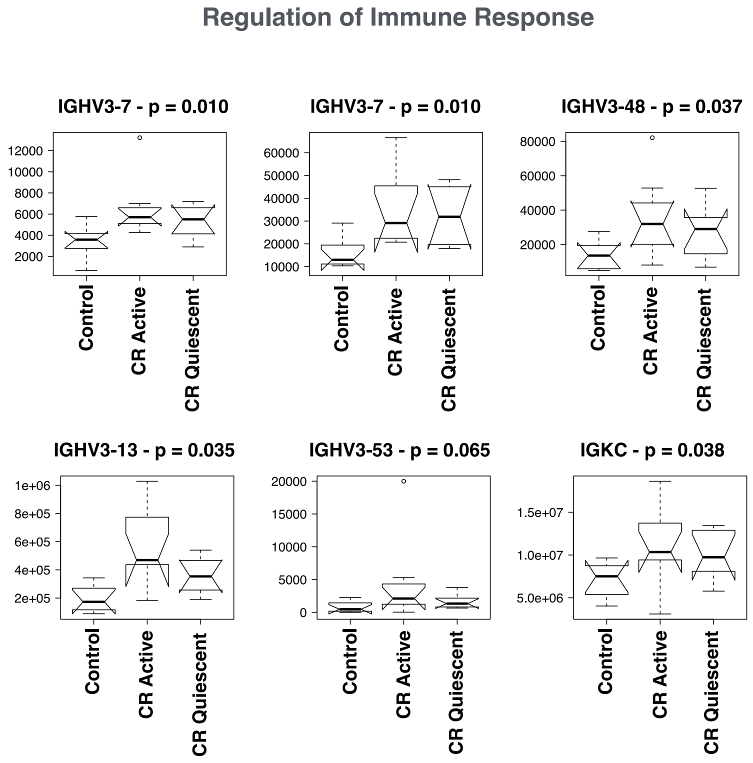

**Figure S5. Similar patterns of protein expression in the CR magenta WGCNA selected module.** The IGHV3 protein family and IGKC barely present the same expression levels between active and quiescent CR status. The second IGHV3-7 is really IGKV2-30.

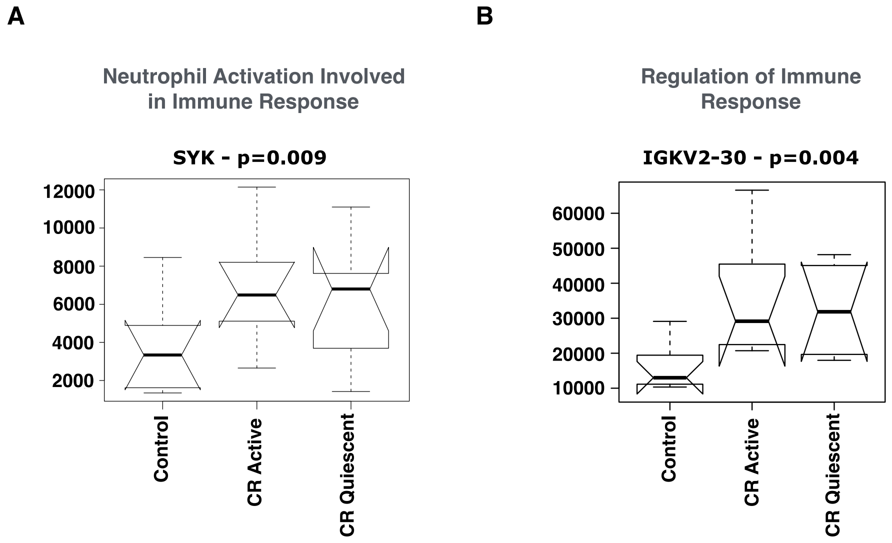

**Figure S6. Patterns of protein expression in the CR magenta WGCNA selected module.** SYK and IGKV2-30 present themselves more expressed in the quiescent than in the active status during CR progression.

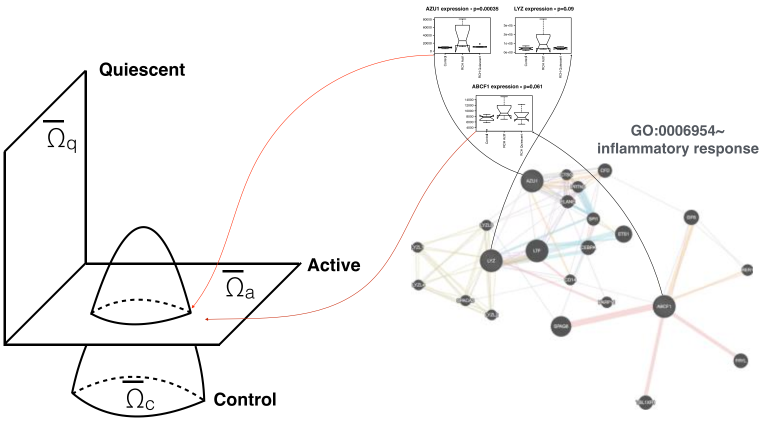

**Figure S7. Abstraction of the IBD progression.** We describe the domain of

IBD progression as intersection of three manifolds eventually with boundaries namely: $\Omega_c$, $\Omega_q$ and $\Omega_a$ to control, quiescent and active status respectively. The data geometry of IBD status can be learnt through the IBD potential introduced in the main manuscript by using in an on top space the differential structure defined by their WGCNA selected proteins.

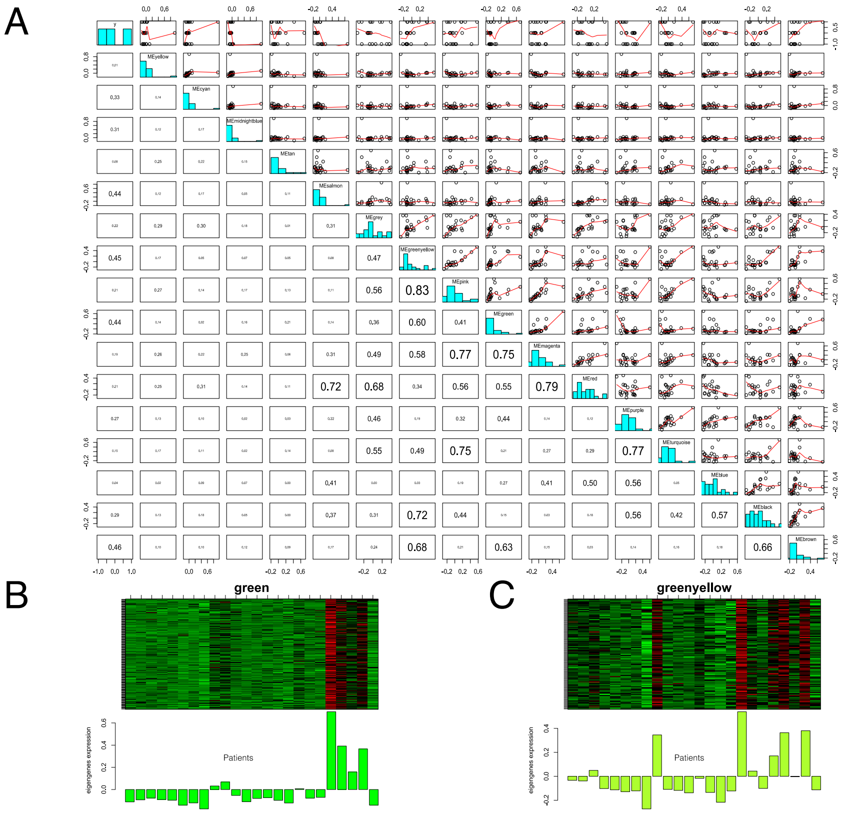

**Figure S8. Estimate functions generated by the eigengenes in UC. (A)** Dependency boxplots for every correlated module in UC. **(B)** Eigengenes of the green module. **(C)** Eigengenes of the greenyellow module.

A

B

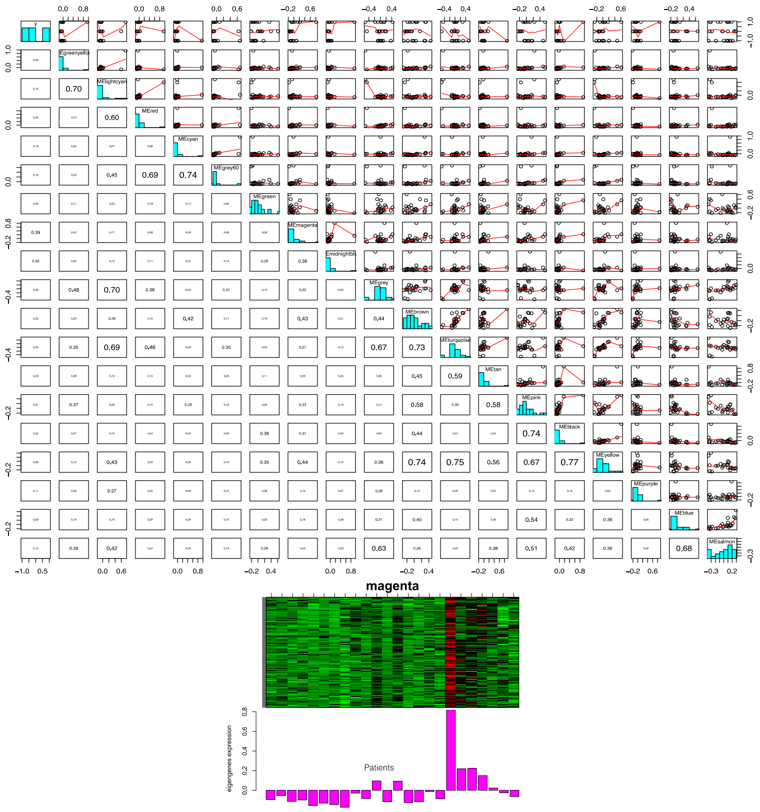

**Figure S9. Estimate functions generated by the eigengenes in CR. (A)** Dependency boxplots for every correlated module in CR. **(B**) Eigengenes of the magenta module.

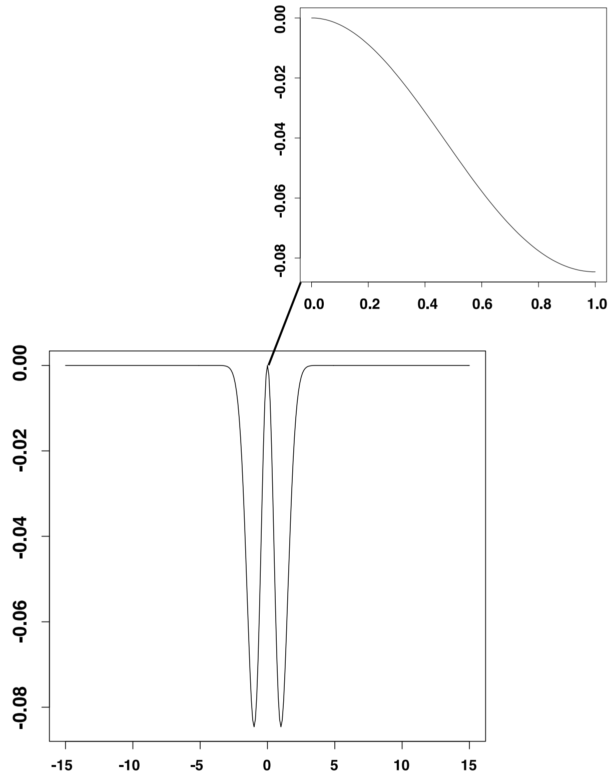

**Figure S10. Potential of the UC/CR control-active set.** The geometry described in the principal manuscript, coupled with the abstracted disease-related dynamics through this potential defined by the equation $L_h f(x) = \frac{1}{\sqrt{h}} C r e^{-r^2}$, can be used to prioritise therapeutic interventions. Inset: zoom into the positive unitary domain.
